# MLL4 regulates postnatal palate growth and midpalatal suture development

**DOI:** 10.1101/2024.07.16.603832

**Authors:** Jung-Mi Lee, Hunmin Jung, Bruno de Paula Machado Pasqua, Yungki Park, Qinghuang Tang, Shin Jeon, Soo-Kyung Lee, Jae W. Lee, Hyuk-Jae Edward Kwon

**Author notes:** Corresponding authors: Hyuk-Jae Edward Kwon, D.D.S., Ph.D., Department of Oral Biology, School of Dental Medicine, University at Buffalo, The State University of New York, 547 Biomedical Research Building, Buffalo, NY 14214-8024, U.S.A., Jung-Mi Lee, Ph.D., Department of Oral Biology, School of Dental Medicine, University at Buffalo, The State University of New York, 540 Biomedical Research Building, Buffalo, NY 14214-8024, U.S.A.

## Abstract

MLL4, also known as KMT2D, is a histone methyltransferase that acts as an important epigenetic regulator in various organogenesis programs. Mutations in the *MLL4* gene are the major cause of Kabuki syndrome, a human developmental disorder that involves craniofacial birth defects, including anomalies in the palate. This study aimed to investigate the role of MLL4 and the underlying mechanisms in the development and growth of the palate. We generated a novel conditional knockout (cKO) mouse model with tissue-specific deletion of *Mll4* in the palatal mesenchyme. Using micro-computed tomography (CT), histological analysis, cell mechanism assays, and gene expression profiling, we examined palate development and growth in the *Mll4*-cKO mice. Gross craniofacial examination at adult stages revealed mild midfacial hypoplasia and midline defects of the palate in *Mll4*-cKO mice, including a widened midpalatal suture and disrupted midline rugae pattern. Micro-CT-based time-course skeletal analysis during postnatal palatogenesis through adulthood demonstrated a transverse growth deficit in overall palate width in *Mll4*-cKO mice. Whole-mount and histological staining at perinatal stages identified that the midline defects in the *Mll4*-cKO mice emerged as early as one day prior to birth, presenting as a widened midpalatal suture, accompanied by increased cell apoptosis in the suture mesenchyme. Genome-wide mRNA expression analysis of the midpalatal suture tissue revealed that MLL4 is essential for the timely expression of major cartilage development genes, such as *Col2a1* and *Acan*, at birth. Immunofluorescence staining for osteochondral differentiation markers demonstrated a marked decrease in the chondrogenic marker COL2A1, while the expression of the osteogenic marker RUNX2 remained unchanged, in the *Mll4*-cKO midpalatal suture. Additionally, SOX9, a master regulator of chondrogenesis, exhibited a significant decrease in protein expression. Indeed, time-course histological analysis during postnatal palate growth revealed retardation in the development of the suture cartilage in *Mll4*-cKO mice. Taken together, our results demonstrate that MLL4 is essential for orchestrating key cellular and molecular events that ensure proper midpalatal suture development and palate growth.

## Introduction

The palate is a central structure in the craniofacial complex, playing a crucial role in breathing, feeding, speech, and maintaining overall craniofacial health. Thus, palatogenesis—the development and growth of the palate—is a vital process in establishing the craniofacial complex. Palatogenesis can be broadly divided into two phases: embryonic palatogenesis and postnatal palatogenesis, which occurs very similarly in humans and mice (Dixon et al., 2011). The first phase, embryonic palatogenesis, occurs during the early stages of fetal development and involves the initial formation of a pair of palatal shelves, which are bilateral outgrowths from the left and right maxillary processes, at embryonic day (E) 11 in mice. These bilateral palatal shelves then undergo remodeling, grow toward the midline, and fuse with each other between E12 and E15, establishing the foundational structure of the palate (Bush and Jiang, 2012). The second phase, postnatal palatogenesis, involves skeletal growth, which occurs along the anteroposterior and transverse directions, with the palate reaching its full adult dimension by 5 weeks of age in mice. A key feature in postnatal palatogenesis is the role of the palatal sutures, which remain as active growth sites postnatally, allowing for the transverse growth of the palate and the accommodation of the growing craniofacial complex during the growth spurts of childhood and adolescence (Griffiths et al., 1967, Latham, 1971). For example, infant patients that undergo surgical repair of cleft palate frequently exhibit signs of midfacial growth arrest, whereas those with unrepaired cleft palate often display normal jaw dimensions and occlusion (Fudalej et al., 2008, Mølsted et al., 2005, Brattström et al., 2005, Wolford et al., 2008, Ye et al., 2012). Similarly, in developing mice, surgical disruption of the midpalatal suture can result in a narrow palate and midfacial hypoplasia (Li et al., 2015). Both phases are crucial for the functional integrity of the palate, and their regulation is orchestrated by a complex interplay of genetic and epigenetic factors, ensuring the precise expression of developmental genes at the right time and place (Li et al., 2017). Any disruption in these factors may lead to severe craniofacial birth defects such as orofacial clefts and midfacial hypoplasia (Garland et al., 2020, Reynolds et al., 2020, Cox, 2004, Suzuki et al., 2016). This direct clinical relevance has driven substantial research efforts to understand embryonic palatogenesis. In contrast, postnatal palatogenesis remains significantly understudied, especially regarding the underlying mechanisms, which has contributed to disparity in research focus. Thus, there is a critical need to expand research into postnatal palatogenesis to address issues related to functional impairments and orthodontic concerns that can arise later in life and to better understand the complete picture of craniofacial development.

Epigenetic regulation, including histone modification, plays a pivotal role in orchestrating craniofacial development and growth (Shull and Artinger, 2024). By regulating chromatin structure and accessibility, epigenetic regulation ensures precise temporal and spatial expression of genes essential for establishing the craniofacial structures in health and disease, influencing cellular mechanisms, such as proliferation, migration, and differentiation. In this regard, mixed lineage leukemia 4 (MLL4), also known as histone-lysine *N*-methyltransferase 2D (KMT2D), is a plausible candidate that may serve as a key epigenetic regulator in palatogenesis. MLL4 is a histone H3 lysine 4 (H3K4) methyltransferase that plays a crucial role in modulating gene expression patterns that are fundamental to the establishment and maintenance of cell identity and lineage specificity during organogenesis (Froimchuk et al., 2017). Mutations in *MLL4* cause Kabuki syndrome, which is a human developmental disorder characterized by distinctive facial features including eversion of lower lateral eyelids and elongated palpebral fissures, postnatal growth retardation, intellectual disability, and skeletal, craniofacial, and dermatoglyphic abnormalities (Boniel et al., 2021). Within the craniofacial complex, Kabuki syndrome is associated with orofacial anomalies, including various palate anomalies and midfacial dysplasia (Matsune et al., 2001). Previous studies using various animal models have shown that MLL4 plays an important role in craniofacial development by regulating neural crest formation, migration, and differentiation (Bögershausen et al., 2015, Schwenty-Lara et al., 2020, Van Laarhoven et al., 2015). Notably, mice with tissue-specific deletion of *Mll4* in the neural crest cell-lineage, driven by either *Wnt1-Cre* or *Sox10-Cre*, showed cleft palate (Shpargel et al., 2020). This suggests that palatal shelf fusion during embryonic palatogenesis requires MLL4 function in the cranial neural crest-derived cells, which give rise to the palate mesenchyme.

However, deletion of *Mll4* in these mice occurs considerably earlier than the initiation of palatogenesis (at E11). Therefore, it remains uncertain whether the cleft palate phenotype is a result of *Mll4* deficiency during or prior to the initiation of palatogenesis. Moreover, due to the neonatal lethality, these neural crest-specific *Mll4* deletion mice were limited to investigating only the embryonic palatogenesis. Thus, there is a critical need to understand the comprehensive role and underlying mechanism of MLL4 in palatogenesis in a time- and tissue-specific manner.

In this study, to elucidate the comprehensive role and the underlying mechanism by which MLL4 regulates palatogenesis, we generated a novel mouse model with palate mesenchyme-specific and palatogenesis-specific *Mll4* deletion (*Mll4^fl/fl^;Osr2-Cre*). Using micro-computed tomography (CT), histology, cell mechanism assays, and gene expression analysis approaches, we examined the development and growth of the palate in our mouse model. Our results revealed that *Mll4* deficiency results in retardation in the developmental of the midpalatal suture and transverse growth deficits during postnatal palatogenesis. Our study provides critical insights into the intrinsic role of MLL4 in palatogenesis, which informs the physiological developmental mechanism and suggests therapeutic strategies for congenital orofacial anomalies.

## Materials and Methods

### Mouse Strains

*Osr2-Cre* (originally described as ‘*Osr2KI*-Cre’) (Chen et al., 2009, Shpargel et al., 2020), and *Mll4^fl/fl^* (Lee et al., 2013) lines were used for generating an *Mll4* conditional knockout (cKO) model. *ROSA^mT/mG^* (hereafter referred to as *mTmG*) reporter line (Muzumdar et al., 2007) was used to label palate mesenchyme cells expressing eGFP by crossing with *Osr2-Cre* line. Mice were maintained by intercrossing or by crossing with C57BL/6J inbred mice. All animal procedures were approved by the Institutional Animal Care and Use Committee at the University at Buffalo.

### Micro-CT Scanning and 3-Dimensional Reconstruction

The heads of mice aged 1, 3, 6, and 10 weeks were fixed in 10% formalin overnight and subsequently washed with PBS. Micro-CT scans were performed using a SCANCO µCT 100 (SCANCO Medical, Switzerland) (voxel size, 10 µm). The scanned images were calibrated to milligrams of hydroxyapatite per cubic centimeter (mgHA/cm^3^) for accurate density measurement. Each image was manually adjusted for rotation and alignment along each axis using the FIJI software (Schindelin et al., 2012). To eliminate background noise, areas with measurements below 200.03 mgHA/cm^3^ were removed. Each image was then cropped to focus on the specific region of interest within the maxillary jaw. The Fijiyama plugin was used to manually register the images, aligning them for further analysis (Fernandez and Moisy, 2021). Finally, the 3Dscript plugin was used for 3-dimensional (3D) rendering (Schmid et al., 2019).

### Histology and Immunofluorescence Staining

Mice were euthanized at predetermined stages. Head tissue specimens were fixed in 4% paraformaldehyde (PFA), dehydrated through an ethanol series, embedded in paraffin, and sectioned at 5–7 μm thickness, as described previously (Lee et al., 2022). Specimens harvested at E18.5, postnatal day (P) 0.5, and 1, 3, 6, and 10 weeks were decalcified in 14% ethylenediaminetetraacetic acid (EDTA) for 1, 3, 6, and 10 weeks, respectively. For histological analysis, sections were stained with Azan’s trichrome staining, as previously described (Lee et al., 2022), and were imaged on an Axioscope compound upright light microscope (for 20X and 40X magnifications) (Zeiss, Germany). An SZX16 stereomicroscope (Olympus, Japan) was used for 5X magnification images, as well as gross images. For immunofluorescence staining, antigen retrieval was performed in an antigen retrieval buffer, using a pressure cooker for 15 min at low pressure, followed by washing in PBS, blocking at room temperature in a blocking buffer (2% goat serum, 5% BSA, 1% Triton-X 100, 0.1% Tween-20 in PBS) for 1 hr, and incubation with primary antibodies, diluted in blocking buffer, overnight at 4°C. On the second day, the sections were washed in PBS, incubated with secondary antibodies, diluted in blocking buffer, at room temperature for 1 hr, washed in PBS, and mounted with Vectashield mounting media with DAPI (Vector Laboratories, USA; H-1200). The primary antibodies used were: homemade guinea pig anti-MLL4 (1:300) (Huisman et al., 2021), rabbit monoclonal anti-SOX9 (1:300; Abcam, UK; ab185230), mouse monoclonal anti-COL2A1 (1:50, Santa Cruz, USA; sc-52658), and rabbit monoclonal anti-RUNX2 (1:300; Cell Signaling, USA; #12556). Immunofluorescence images were acquired using a THUNDER Imager fluorescence microscope (Leica Microsystems, Germany). To eliminate non-specific signals from auto-fluorescence, we acquired additional images usng a non-stained CFP (cyan) filter and subtracted these signals following an established scientific method (Van de Lest et al., 1995).

### TUNEL Assay

The DeadEnd™ Fluorometric TUNEL System (Promega, USA; G3250) was used to assess apoptosis in paraffin sections, according to the manufacturer’s protocol.

### Skeletal Preparations

Mouse heads, harvested at predetermined stages, were fixed in 100% ethanol for 3 days, stained in Alcian Blue solution (30 mg Alcian Blue dissolved in 20 ml glacial acetic acid plus 80 ml 95% ethanol) for 2 days, re-fixed in 100% ethanol for 24 hr, cleared in 2% KOH for 3 hr, and counterstained in Alizarin Red solution (50 mg Alizarin Red dissolved in 1 L of 1% KOH) for 24 hr, followed by tissue clearing in 1% KOH/20% glycerol for 3 days. Images were acquired using a standard stereomicroscope (Amscope, USA).

### RNA Sequencing and Bioinformatics Analysis

Total RNA was extracted from micro-dissected midpalatal suture tissue samples using the RNeasy Micro Plus Kit (Qiagen, Germany; 74034). Four biological replicates per genotype (control and *Mll4*-cKO), were used for sequencing, with one sample in each genotype pooled from three individuals. Library construction and high-throughput sequencing were performed by Novogene company (China) using the Illumina NovaSeq 6000 platform. Sequencing reads were analyzed using Salmon (Patro et al., 2017) to quantify gene expression at the transcript level. The resulting outputs from Salmon were analyzed using DESeq2 (Love et al., 2014) to obtain gene-level expression data and identify differentially expressed genes (DEGs) between *Mll4*-cKO and littermate control samples. DEGs were defined by a log2 fold change (log2FC) ≥ 0.4 and an adjusted *p*-value ≤ 0.001. Volcano plots were generated using the EnhancedVolcano package (Blighe et al., 2024), and gene ontology (GO) analysis was performed with clusterProfiler (Wu et al., 2021). Raw expression data results are included in the Supplementary Material. The original RNA sequencing (RNA-seq) data files have been deposited in the National Center for Biotechnology Information Gene Expression Omnibus (NCBI GEO) database under the accession number PRJNA1148613.

### ChIP Sequencing Re-Analysis

The ChIP-sequencing (ChIP-seq) datasets for UTX in cranial neural crest cells (GSE103849) were re-analyzed using a previously described data processing workflow (Shpargel et al., 2017). Briefly, sequencing reads were aligned to the mouse mm10 reference genome using Bowtie2 (Langmead and Salzberg, 2012) and converted to BAM files using SAMtools (Li et al., 2009). Reads mapping to the X and Y chromosomes were excluded. Filtered BAM files from biological replicates were merged and subjected to peak calling with MACS (Zhang et al., 2008). Peaks were annotated using the ChIPseeker R package (Yu et al., 2015). For visualization in Integrative Genomics Viewer (IGV), the filtered BAM files were further processed using BEDTools (Quinlan and Hall, 2010) and converted to BigWig format using the UCSC command-line tool bedGraphToBigWig (Kent et al., 2002). The intersection of *Mll4*-cKO DEGs with UTX target genes, along with functional enrichment analysis, was performed in R using clusterProfiler (Wu et al., 2021).

### Statistics

All statistical analyses were performed using Prism 10 software (GraphPad Software, USA). Pairwise differential expression was analyzed using Student’s *t*-test. Statistical significance was defined as *p*-values ≤ 0.05 (*), ≤ 0.01 (**), and ≤ 0.001 (***).

## Results

### Generating a novel *Mll4* deficiency model for studying the palate tissue-autonomous role of MLL4 during palate development and growth

For the purpose of investigating the role of MLL4 specifically during palatogenesis, we used the Cre/loxP system driven by *Osr2-Cre*. *Osr2-Cre* is first activated at E11, when palatogenesis is initiated, and remains active in the palate mesenchyme cells (Lan et al., 2001, Lan et al., 2004). The *Osr2-Cre;mTmG* line allows us to confirm the activation of *Osr2-Cre* by detecting GFP signals using fluorescence microscopy. When the bilateral palatal shelves are actively undergoing morphogenesis, for example at E13.5, *Osr2-Cre* was activated in the developing palatal shelves, which occupies the center of the developing face (Fig. 1A, B). Thus, we were able to generate a novel mouse model with palatogenesis-specific deletion of *Mll4* in the palate mesenchyme cells (*Mll4^fl/fl^;Osr2-Cre*, or *Mll4*-cKO). Surprisingly, compared to the previously reported neural crest-specific *Mll4* deletion (*Mll4^fl/fl^;Wnt1-Cre* and *Mll4^fl/fl^;Sox10-Cre*) mice, which exhibited cleft palate and neonatal lethality (Shpargel et al., 2020), our palatogenesis-specific *Mll4*-cKO mice did not show cleft palate or lethality at birth and could grow up into adult mice that did not show much noticeable changes when grossly examined for exterior appearance at 10 weeks (Fig. 1C, G). However, close examination of the orofacial skeleton dimensions by using micro-CT analysis revealed that the facial angle, length, and width in the maxilla showed mild but significant decreases in the *Mll4*-cKO mice (Fig. 1D–F, H–J, N), which were signs of mild midfacial hypoplasia. Additional examination of the intra-oral aspect of the palate revealed several minor differences between the control and *Mll4*-cKO mice (Fig. 1K, L). Compared to controls, the midpalatal suture appeared more widened in the *Mll4*-cKO palate (Fig. 1K′, L′, arrowheads), with minor changes observed in the rugae patterning (Fig. 1K′, L′, arrows). To test the possibility that these mild midfacial hypoplasia and palate phenotypes that we observed in the *Mll4*-cKO mice may have resulted from delay in their overall development and growth, we measured the body weight both at 10 weeks and at birth but did not find any significant difference (Fig. 1M). We also compared the body weight between male and female mice at these two stages but found no significant difference (data not shown). Our initial observations suggested that, separate from its early role in neural crest cell differentiation, MLL4 may have other unknown functions in postnatal palate development and growth.

**Figure 1.**
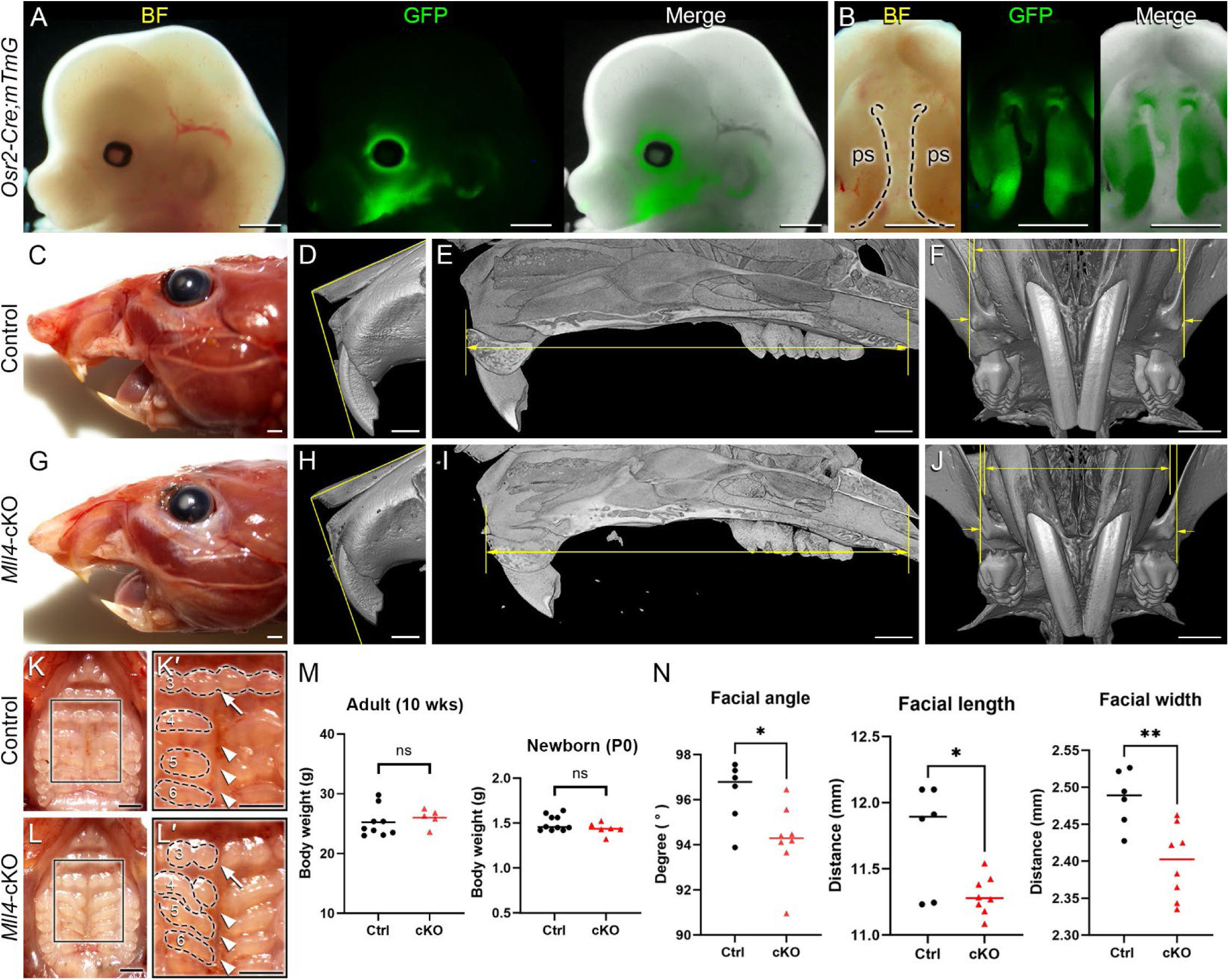
Generation of a novel *Mll4*-deficient Kabuki syndrome mouse model for palate studies. A conditional *Mll4*-deficiency mouse model (*Mll4^fl/fl^;Osr2-Cre*, or *Mll4*-cKO) was generated, using the Cre/loxP system driven by *Osr2-Cre* for studying postnatal palate development. (A, B) Osr2-Cre activation in the developing midface (A) and palate (B), demonstrated in the *Osr2-Cre;mTmG* mouse at E13.5. ps, palatal shelf (in B). (C–J) Craniofacial phenotype analysis, including gross examination (C, G) and micro-CT analysis (D–F; H–J): facial angle (D, H), facial length (E, I), and facial width (F, J) all show mild but significant decrease in the *Mll4*-cKO (H–J) compared to control (D–F) in adult mice at 10 weeks. (K, L) Orofacial phenotypes in the control (K, K′) and *Mll4*-cKO (L, L′) adult mice at 10 weeks: *Mll4*-cKO mice exhibit midline defects, such as widened midpalatal suture (arrowheads) and discontinuous ruga (3, arrows), along with disrupted rugae patterning (3– 6). (M) Body weight comparison between the control and *Mll4*-cKO mice shows no significant difference in either adult (*n*=9 and 5 for control and cKO, respectively) or newborn mice (*n*=10 and 6 for control and cKO, respectively). (N) Measurements of facial angle, length, and width from panels D–F and H–J (*n*=6 for both control and cKO). Scale bars: 1 mm (in panels A–L).

Thus, to gain better understanding of the midfacial and palate phenotypes observed from gross examination of the *Mll4*-cKO mice (Fig. 1), we further examined the palate skeleton at 10 weeks using our micro-CT results. By examining the virtual 3D models that were created by reconstructing the planar X-ray images, we found that the overall palate width showed narrowing in the *Mll4*-cKO mice, which was consistent with the decrease in the facial width that we observed earlier. Anatomical landmarks for assessing the palate width, such as the incisive foramens, palatal processes of maxilla, and palatal processes of palatine, all showed constricted appearances in the virtual 3D model of the *Mll4*-cKO mice (Fig. 2A, B). Using virtual slicing of the 3D models, we were able to further generate digital coronal/frontal cross sections of the palate. These virtual cross sections enabled us to perform comprehensive quantitative analysis of the overall horizontal dimension of the palate along its anteroposterior axis by precisely measuring the palate width at four representative section levels: the incisive foramen (in slice 1), the palatal processes of maxilla (in slice 2), the transpalatal suture (in slice 3), and the palatal processes of palatine (in slice 4) (Fig. 2C, D). At all four section levels along the anteroposterior axis, the palate width showed significant decrease in the *Mll4*-cKO mice, compared to the control mice (Fig. 2E1–E4). These results suggested that MLL4 plays an important role in transverse growth of the palate during postnatal palatogenesis.

**Figure 2.**
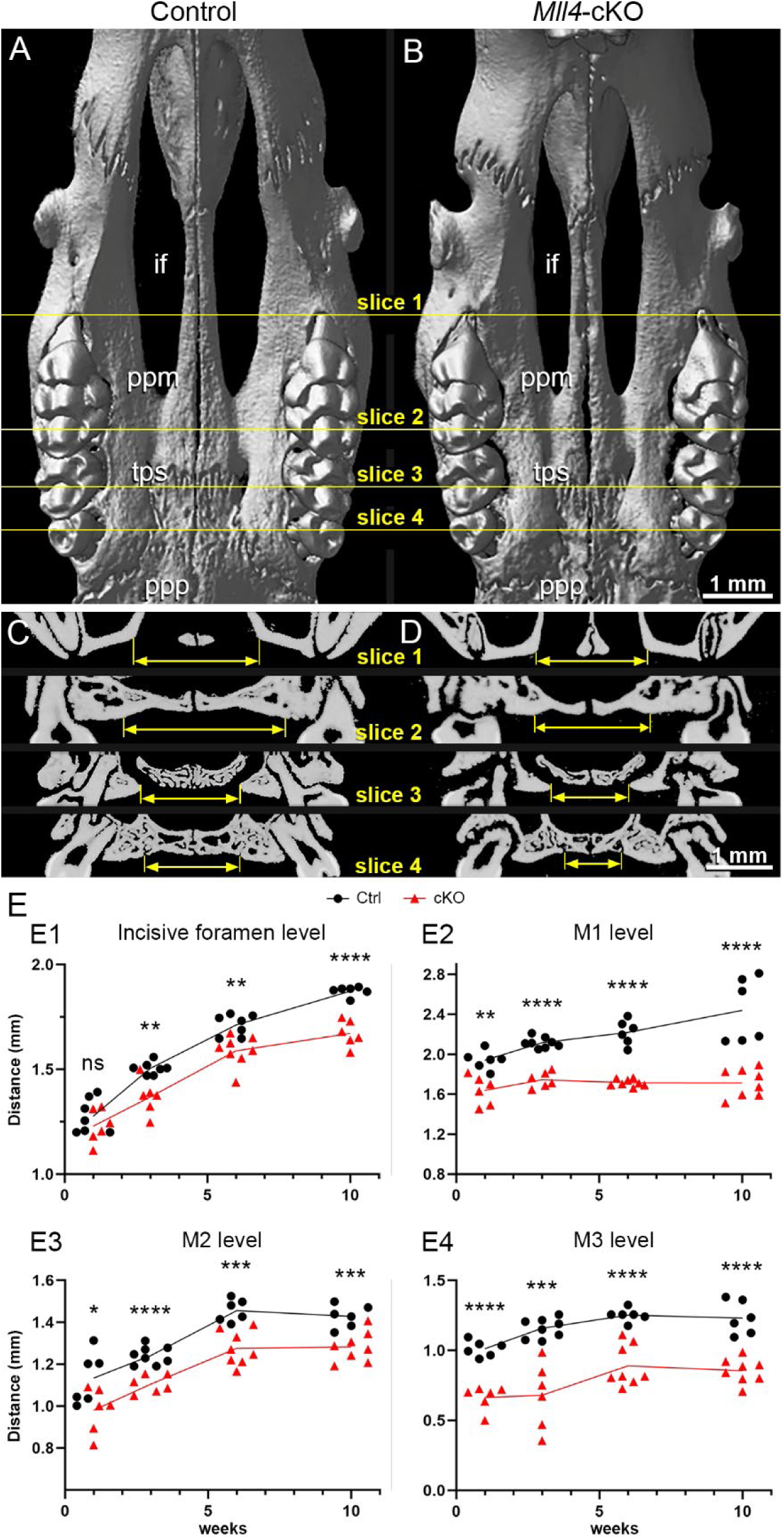
*Mll4* is essential for proper growth of the palate during postnatal palatogenesis. (A–D) Skeletal phenotypes were examined, using micro-CT analysis, in control (A, C) and *Mll4*-cKO (B, D) adult mice at 10 weeks. Virtual 3D (A, B) and coronal/frontal sections (C, D) along the anteroposterior axis at four representative levels—for the incisive foramen (if) (slice 1), the palatine process of maxilla (ppm) (slice 2), transpalatal suture (tps) (slice 3), and the palatal process of the palatine (ppp) (slice 4)—were examined. (E) Palate width was measured in each for the four sections that represent the incisive foramen (slice 1), the palatine process of maxilla (slice 2), transpalatal suture (slice 3), and the palatal process of the palatine (slice 4) at 1, 3, 6, and 10 weeks. *n*=6 or more for both controls and cKOs at all stages. Scale bars: 1 mm (in panels A–D).

Thus, to address the question of whether MLL4 is actually required for proper growth of the palate during postnatal palatogenesis, we conducted a time-course quantitative analysis of the horizontal dimension of the palate skeleton using micro-CT-generated virtual coronal/frontal cross sections at 1, 3, and 6 weeks. For this approach, we used the same four representative coronal/frontal cross section levels along the anteroposterior axis of the palate, which we used at 10 weeks. At 1 week, we found that palate growth was not significantly changed in the *Mll4*-cKO mice at the incisive foramen level (Fig. 2E1). In contrast, at the rest of the levels, the first (M1), second (M2), and third molar (M3) levels, palate growth showed minor, but significant decrease (Fig. 2E2–E4). Later, at 3, 6, and 10 weeks, palate growth showed significant decrease at all four levels along the anteroposterior axis in the *Mll4*-cKO mice, compared to the control mice (Fig. 2E1–E4). These results suggested that MLL4 is indeed required for proper transverse growth of the palate during postnatal palatogenesis.

### MLL4 is essential for the initial formation of the midpalatal suture around birth

We next asked the question of how MLL4 promotes palate growth. Based on our finding that the *Mll4*-cKO mice showed significant narrowing of the palate during postnatal palatogenesis (Fig. 2), we reasoned that the key biological events for palate growth that depend on MLL4 function may occur at younger ages, possibly as early as, or even prior to, birth. Thus, to identify the primary developmental mechanism underlying the role of MLL4 in palate growth, we examined the palate anatomy in the control and *Mll4*-cKO mice around birth, searching for the earliest signs of disruption in the *Mll4*-cKO palate. By first using skeletal staining of the developing cranium and palate tissues, we found that the overall skeletal pattern was similar between the control and *Mll4*-cKO mice (Fig. 3A, B). However, where we had initially observed midline defects in the rugae pattern and a widened midpalatal suture in the *Mll4*-cKO mice (Fig. 1), we found that the width of the midpalatal suture was significantly wider in the *Mll4*-cKO mice, compared to the controls, prior to birth (at E18.5) (Fig. 3A, B, G). In parallel, histological examination of the palate, by using trichrome staining of the coronal/frontal paraffin sections, confirmed that the width of the midpalatal suture, situated between the bilateral bony palatal processes, was significantly increased in the *Mll4*-cKO mice, compared to the control mice (Fig. 3C, D, C′, D′, G). To determine the cellular mechanisms underlying the wider midpalatal suture in the *Mll4*-cKO mice, we performed both cell proliferation, using EdU incorporation assays, and cell apoptosis, using TUNEL staining. While rarely any cells in the midpalatal suture were EdU-positive for cell proliferation in both the control and *Mll4*-cKO mice (data not shown), TUNEL-positive cell apoptosis showed minor but significant increase in the *Mll4*-cKO midpalatal suture, compared to controls, at E18.5 (Fig. 3E, F, E′, F′, H). Our findings that the earliest signs of disrupted biological events were specifically discovered in the midpalatal suture of the perinatal *Mll4*-cKO palate suggested that MLL4 is essential for the establishment of the midpalatal suture around birth.

**Figure 3.**
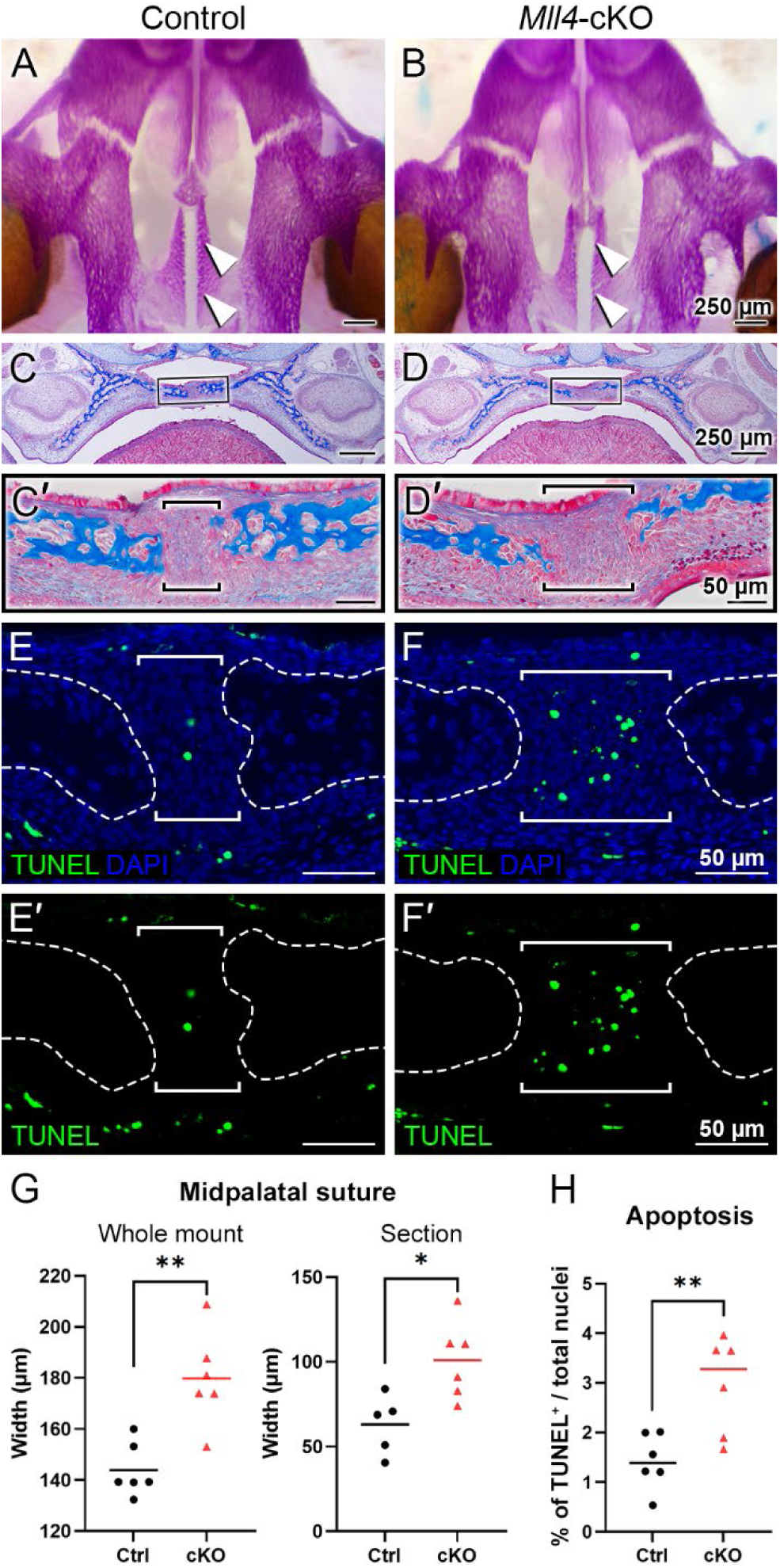
*Mll4* is required for initial formation of the midpalatal suture prior to birth. (A–D, G) Skeletal staining (A, B) and trichrome staining results (C, D), with quantitative analysis of the midpalatal suture width (G). Bone is stained magenta (in A, B) or blue (in C, D). The overall skeletal patterning was similar between the control (A, C) and *Mll4*-cKO palate (B, D) prior to birth (at E18.5). However, specifically in the newly established midpalatal suture (arrowheads in A, B; brackets in C′, D′), *Mll4*-cKO mice (D, D′) showed wider suture width, compared to the control mice (C, C′, G). (E, F, E′, F′, H) TUNEL staining at E18.5 revealed that *Mll4* deficiency resulted in subtle but significant increase in cell apoptosis in the midpalatal suture (brackets in E, F, E′, F′; H). *n*=6 for both control and cKO in panels G and H.

### *Mll4* deficiency results in dysregulation of timely expression of chondrogenic genes at birth

To gain insight into the molecular mechanism underlying the histological and cellular alterations observed in the perinatal *Mll4*-cKO palate tissue, we micro-dissected the midpalatal suture tissues and performed RNA-seq analysis on control and *Mll4*-cKO mice at birth (P0.5). Our results identified a total of 117 DEGs between the control and *Mll4*-cKO midpalatal suture tissues, with 40 genes upregulated and 77 genes downregulated in the *Mll4*-cKO group (adjusted *p*-value ≤ 0.001, log2FC ≥ 0.4) (Fig. 4A). Notably, among the DEGs were genes associated with cartilage development, or chondrogenesis, including extracellular matrix genes *Col2a1*, *Col9a1*, *Col9a2*, *Acan* (Aggrecan), and *Comp* (Cartilage oligomeric matrix protein), as well as *Mia*/*Cdrap1* (Melanoma inhibitory activity) and *Snorc* (Small novel rich in cartilage). GO analysis identified the top 10 most enriched biological processes, along with a list of downregulated genes associated with each process (Fig. 4B–D). This analysis suggested that cartilage development, endochondral bone morphogenesis, and chondrocyte differentiation are among the highly enriched biological processes potentially regulated by MLL4.

**Figure 4.**
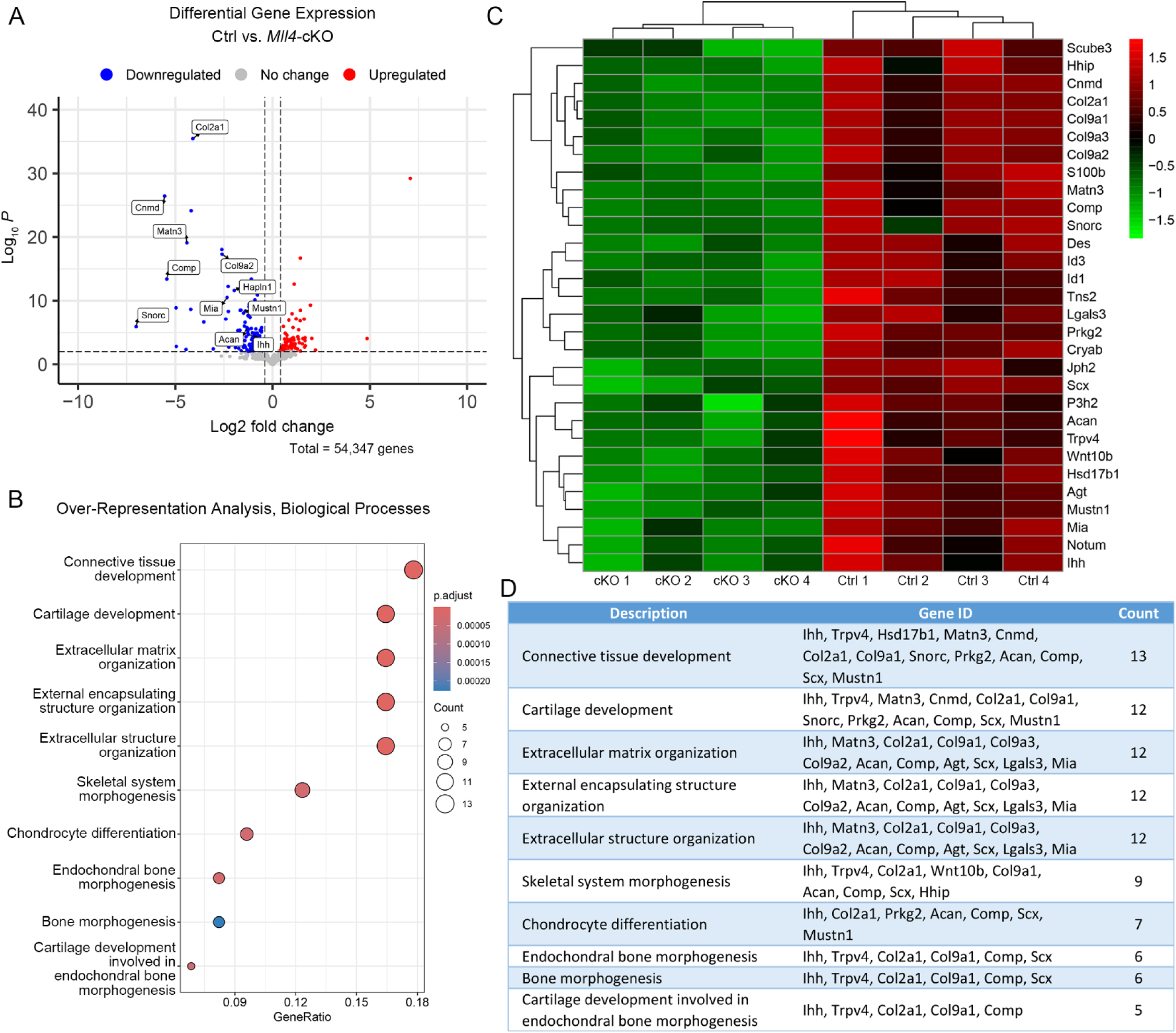
MLL4 regulates expression of genes associated with skeletal development in the newborn midpalatal suture. (A–D) Genome-wide mRNA expression analysis was performed using the control and *Mll4*-cKO midpalatal suture tissues at birth (P0.5). (A) Volcano plot showing that total 117 genes were differentially expressed (adjusted *p*-value ≤ 0.001, log2FC ≥ 0.4)—40 upregulated genes and 77 downregulated genes. (B– D) Top 10-enriched biological processes (B) and heatmap (C), identified through Gene Ontology (GO) analysis from a list of significantly downregulated genes in the *Mll4*-cKO group, compared to the control group (D).

In parallel, we selected several representative markers of osteochondral progenitor cell differentiation during chondrogenesis and endochondral ossification for immunofluorescence staining on paraffin sections of the perinatal palate (at P0.5) in control and *Mll4*-cKO mice. Among these markers, COL2A1 was selected from the DEG list (Fig. 4) to validate our RNA-seq results. Additionally, RUNX2 and SOX9 were chosen as critical transcription factors regulating osteochondral progenitor cell differentiation: RUNX2 marks osteoblast differentiation, while SOX9 marks chondrocyte differentiation, providing insights into the developmental processes occurring in the midpalatal suture. We also verified the expression of MLL4 in the perinatal *Mll4*-cKO palate using immunofluorescence staining. Our results revealed that the expression of COL2A1 and SOX9, both early markers of chondroblast differentiation, was markedly reduced in the *Mll4*-cKO midpalatal suture (Fig. 5B1, D2, E), compared to controls (Fig. 5A1, C2, E). In contrast, the expression of RUNX2, an osteogenic differentiation marker, was similar between the control and *Mll4*-cKO midpalatal suture (Fig. 5A2, B2, E). These findings suggest that the primary molecular events dependent on MLL4 function in the perinatal midpalatal suture are associated with chondrogenic cell differentiation, cartilage development, and potentially endochondral ossification. Additionally, the markedly reduced intensity of MLL4 in the perinatal *Mll4*-cKO palate confirms the effective deletion of the *Mll4* gene in the *Mll4*-cKO palate (Fig. 5C1, D1).

### *Mll4* deficiency leads to retardation of the development of the midpalatal suture

Finally, to verify whether the biological processes that were identified from our RNA-seq analysis were actually impacted as a result of *Mll4* deficiency during postnatal palatogenesis, we performed histological examination of the midpalatal suture at birth and beyond (up to 1 week), using trichrome staining of the paraffin sections (Fig. 6A–D). Interestingly, after birth (at P0.75), a newly formed layer of cartilage tissue capped the medial surface of each of the bilateral palatal processes, flanking the midpalatal suture in the control mice (Fig. 6A, arrowheads). In contrast, the newborn *Mll4*-cKO mice lacked these midpalatal suture-flanking cartilage tissue layers on the medial surfaces of the bilateral palatal processes (Fig. 6B). By histologically monitoring the postnatal development of the midpalatal suture, we found that, at 1 week in the control mice, the midpalatal suture cartilage layers showed uniform enlargement of the consisting chondrocytes (Fig. 6C, arrowheads). In contrast, the 1-week old *Mll4*-cKO mice just started to develop the midpalatal suture cartilage layers, which were only partially formed (Fig. 6D, arrowheads). Our findings suggested that this defective phenotype observed in the *Mll4*-cKO midpalatal suture cartilage was due to significant retardation of chondrogenic differentiation as a result of *Mll4* deficiency in the developing palate. Thus, we further performed additional time-course examination of the midpalatal suture development at 3, 6, and 10 weeks. Interestingly, at all three stages (3, 6, and 10 weeks), the bilateral cartilage layers in both the control and *Mll4*-cKO midpalatal sutures were shown to be adhered to each other in the midline (Fig. 6E–J). However, the interface between the bilateral cartilage layers was less straight in the *Mll4*-cKO suture, compared to the control suture (Fig. 6E–J). Also at all three stages, the overall width of the midpalatal suture cartilage was markedly increased in the *Mll4*-cKO mice (brackets in Fig. 6F, H, J) than in the control mice (brackets in Fig. 6E, G, I). Importantly, compared to the horizontally organized chondrocyte ‘columns’ in the control group, the *Mll4*-cKO suture chondrocytes appeared disorganized and were under-differentiated, especially at 10 weeks (Fig. 6I, J, I′, J′). Taken together, these results suggest that *Mll4* deficiency results in retardation in the developmental of the midpalatal suture during postnatal palatogenesis. And by integrating our findings in the developing midpalatal suture (Fig. 6) and in the growing palate skeleton (Fig. 2), our data suggest that *Mll4* is associated with midpalatal suture development and palate growth during postnatal palatogenesis.

## Discussion

### A novel role of MLL4 in determining palate width during postnatal palatogenesis using a newly developed *Mll4* deficiency/Kabuki syndrome mouse model

In this study, we generated for the first time a unique mouse model demonstrating that MLL4 plays an essential role in determining palate width (Fig. 1). By using *Osr2-Cre* to delete *Mll4* tissue-specifically in the palate mesenchyme from the onset of palatogenesis (at E11) (Lan et al., 2001, Lan et al., 2004), we examined the intrinsic role of MLL4 during the active process of palatogenesis (Fig. 2). Unlike previously reported neural crest-specific *Mll4* deletion models driven by *Wnt1-Cre* or *Sox10-Cre*, which exhibit cleft palate and neonatal lethality (Shpargel et al., 2020), our *Mll4*-cKO mice did not show cleft palate or neonatal lethality. This allowed us to extend our examinations beyond prenatal stages. Importantly, the narrow palate phenotype observed in our *Mll4*-cKO mice closely resembles a common craniofacial phenotype of Kabuki syndrome: the high-arched, narrow palate, reported in 72% (64/89) of individuals with Kabuki syndrome (Matsumoto and Niikawa, 2003, Porntaveetus et al., 2018). This prevalence is twice as high as the 35% (68/196) frequency of cleft lip/palate individuals in Kabuki syndrome (Matsumoto and Niikawa, 2003), which was studied using the aforementioned neural crest-specific *Mll4* deletion models (Shpargel et al., 2020). Our findings further emphasize the clinical relevance of the narrow palate phenotype, especially when accompanied by midfacial hypoplasia, which poses potential risks for various dental and orofacial complications, such as crowded or impacted teeth, breathing difficulties, and further midfacial underdevelopment. Taken together, our unique *Mll4*-cKO model not only highlights a significant craniofacial problem in Kabuki syndrome but also sheds light on an understudied yet critical developmental process that warrants urgent attention. These results underscore the multifaceted role of MLL4 in craniofacial development and disease.

### MLL4 is essential for proper development of the midpalatal suture

Here, we establish MLL4 as a crucial factor in midpalatal suture development, as evidenced by the significant developmental delays observed in the *Mll4*-cKO mice beginning in the perinatal stages (Fig. 3) and persisting throughout postnatal palatogenesis into adulthood (Fig. 6). Whole gene expression profiling of midpalatal suture tissues revealed that *Mll4* is required for the timely expression of a large number of genes related to chondrogenesis, including *Col2a1* (Bell et al., 1997, Lefebvre et al., 1997, Ng et al., 1997), *Col9a1* (Genzer and Bridgewater, 2007, Zhang et al., 2003, Zhou et al., 1998), *Col9a2* (Bernard et al., 2003), *Acan* (Sekiya et al., 2000, Han and Lefebvre, 2008), *Comp* (Liu et al., 2007), *Mia*/*Cdrap1* (Xie et al., 1999), and *Snorc* (Jaiswal et al., 2020) (Fig. 4). Interestingly, *Sox9* did not appear in our DEG list, possibly due to (1) variability introduced during tissue micro-dissection, given its small size of the midpalatal suture and the potential for contamination from adjacent tissues, or (2) the relatively low steady-state expression level of *Sox9* compared to other chondrogenic genes (e.g., RPKM values: *Sox9* = 37.6 vs. *Col2a1* = 787.8 and *Acan* = 308.2), which may obscure detectable differences. Nevertheless, immunofluorescence analysis confirmed that SOX9 expression was markedly downregulated in the mesenchyme cells in the *Mll4*-cKO midpalatal suture (Fig. 5), suggesting that MLL4 may indeed regulate *Sox9* expression.

**Figure 5.**
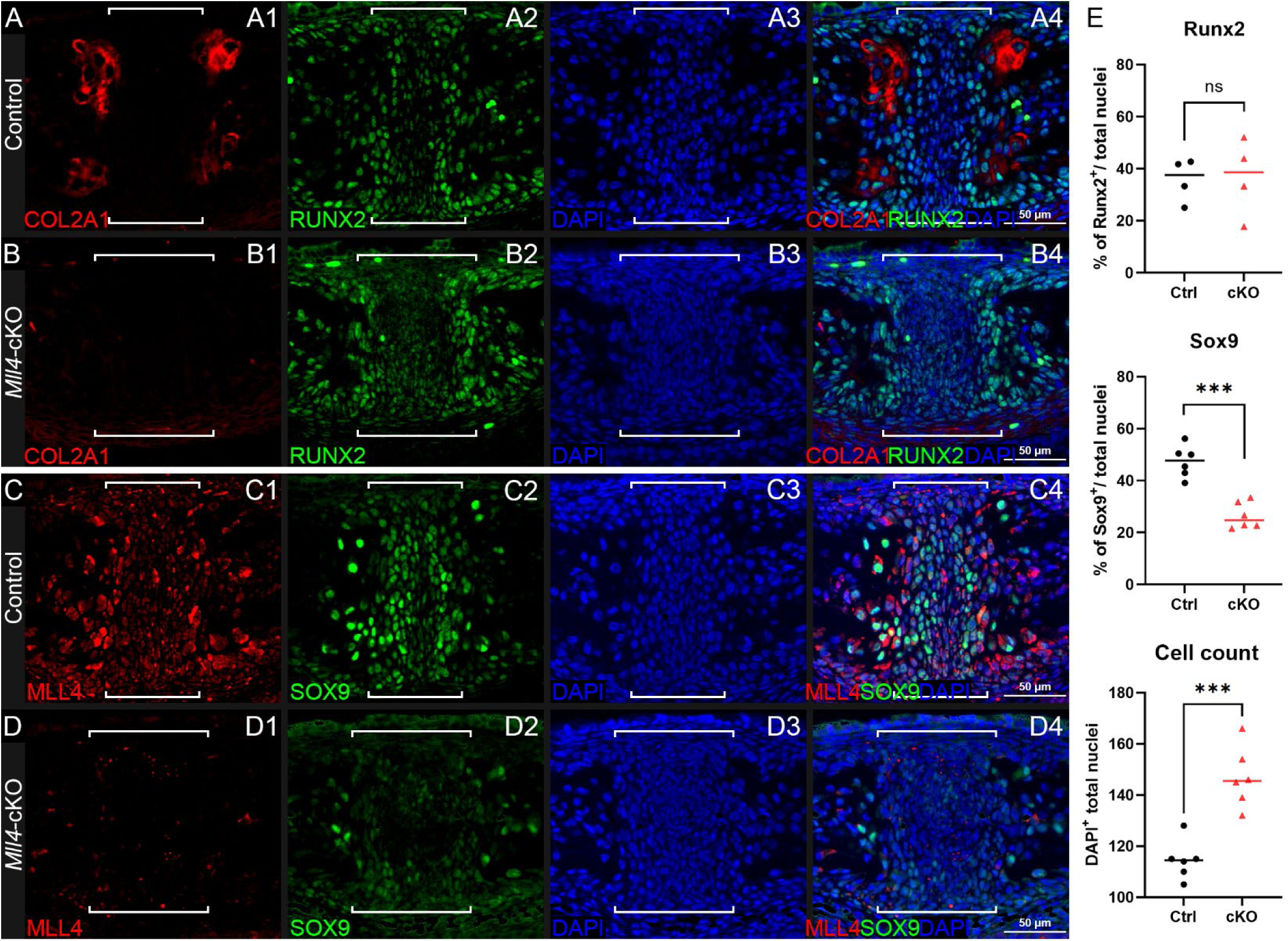
*Mll4* is essential for timely differentiation of chondrogenic cells in the midpalatal suture at birth. (A–D) Immunofluorescence staining for expression of osteochondroprogenitor markers COL2A1, SOX9, and RUNX2, as well as MLL4, in the control (A, C) and *Mll4*-cKO midpalatal suture (B, D) at birth (P0.5): Chondrogenic markers, COL2A1 and SOX9, are markedly reduced in the *Mll4*-cKO midpalatal suture (B1, D2), compared to control (A1, C2). In contrast, osteogenic marker, RUNX2, showed similar expression in the control and *Mll4*-cKO suture tissue (A2, B2). MLL4 was markedly decreased in the *Mll4*-cKO palate (D1), compared to control (C1). Brackets (in A–D) indicate the midpalatal suture. (E) Quantification of RUNX2-positive and SOX9-positive cell ratios, as well as DAPI-positive nuclei counts in the midpalatal suture (*n*=6 for both control and cKO). Scale bars: 50 µm (in panels A–D).

**Figure 6.**
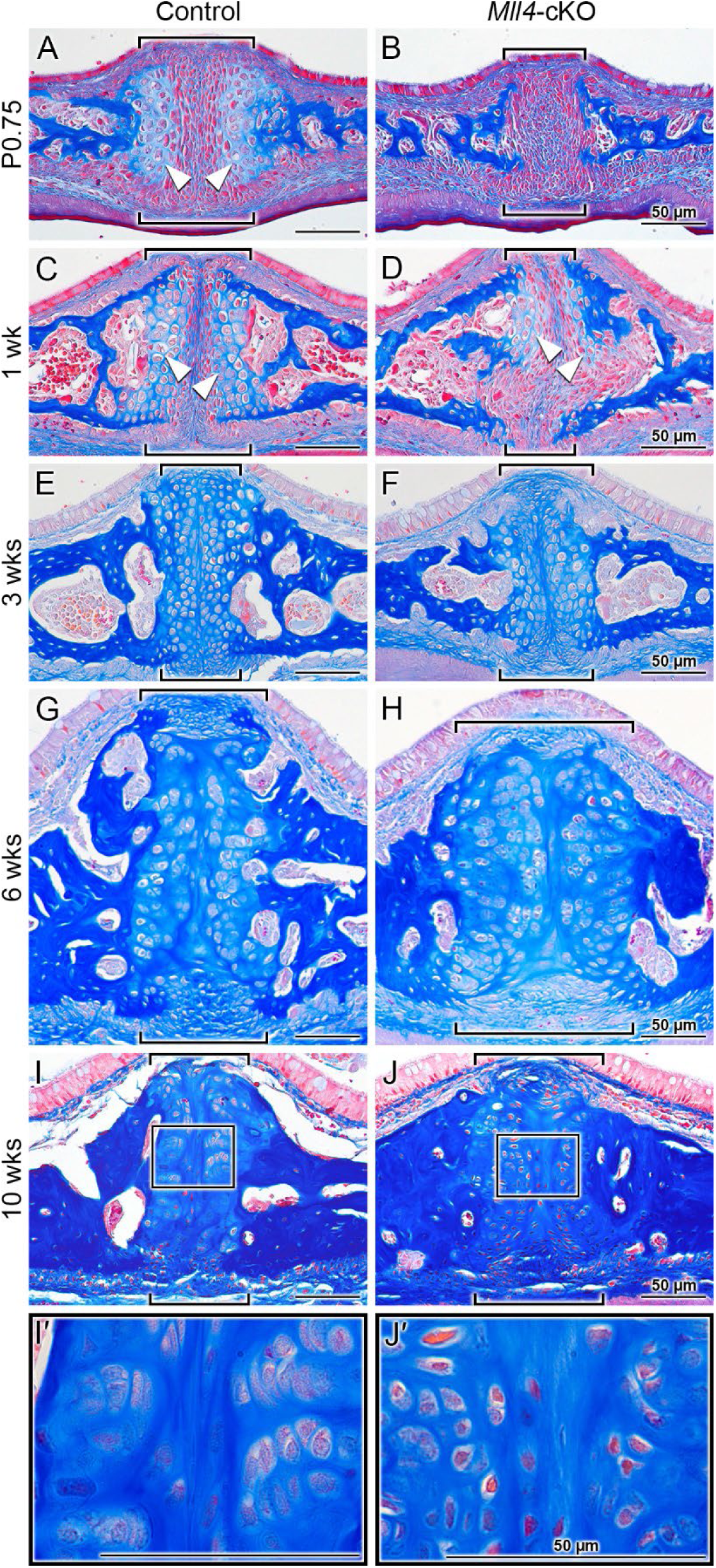
*Mll4* is required for proper development of the midpalatal suture during postnatal palatogenesis. (A–J, I′, J′) Trichrome staining of the midpalatal suture in the control and *Mll4*-cKO mice, showing that *Mll4* deficiency results in significant retardation in the postnatal development of the midpalatal suture cartilage. (A, B) After birth (P0.75), the bilateral cartilage layers become present in the control midpalatal suture (A, arrowheads) but remain completely absent in the *Mll4*-cKO midpalatal suture (B). (C, D) At 1 week, the bilateral midpalatal suture cartilage layers become closer to each other and show uniform enlargement of the chondrocyte cells in the control mice (C, arrowheads), whereas the suture cartilage layers just start to partially appear in the *Mll4*-cKO mice (D, arrowheads). (E–J) At 3 weeks, the bilateral suture cartilage layers adhere to each other in the midline in both the control and *Mll4*-cKO midpalatal sutures, but the *Mll4*-cKO suture (F) is wider than the control suture (E), which becomes more obvious at 6 and 10 weeks (G– J). (I, J, I′, J′) At 10 weeks, the control suture cartilage remains present and narrow (I), and its chondrocytes are well differentiated and organized (I′), whereas the *Mll4*-cKO midpalatal suture cartilage remains wide (J), and its chondrocytes are under-differentiated and disorganized (J′). Brackets (in A–J) indicate the midpalatal suture. Scale bars: 50 µm (in panels A–D).

Notably, the seven downregulated chondrogenic genes in the *Mll4*-cKO midpalatal suture mesenchyme are all known SOX9 targets, highlighting its role as a master regulator of chondrogenesis (Lefebvre et al., 2019). Recent studies suggest that SOX9 functions as a pioneer factor, recruiting epigenetic co-factors such as MLL4 to epigenetically prime and remodel chromatin to a transcriptionally accessible state during cell fate determination (Yang et al., 2023). Therefore, it is reasonable to speculate that *Mll4* deficiency may affect midpalatal suture development through two mechanisms: (1) lack of MLL4 as a transcriptional co-activator of SOX9 and (2) downregulation of SOX9 itself. Both mechanisms would impair SOX9’s ability to activate downstream chondrogenic genes, as observed in the *Mll4*-cKO midpalatal suture. Furthermore, the known auto-regulatory function of SOX9 in chondrocytes (Mead et al., 2013) could exacerbate the chondrogenesis defects in our model.

To further explore the potential mechanism by which MLL4 regulates chondrogenic gene expression, we re-analyzed published UTX ChIP-seq datasets in O9-1 cranial neural crest cells (Shpargel et al., 2017), as MLL4 is known to coordinate with UTX to regulate target gene expression (Cho et al., 2007, Froimchuk et al., 2017). Integrating these datasets with our *Mll4*-cKO DEGs (Fig. 4), we found that approximately 56% (66/117) of the DEGs overlapped with UTX target genes (Fig. 7A), implicating them as potential direct transcriptional targets of MLL4. Functional classification of these genes revealed significant enrichment for skeletal system-related biological processes, including cartilage development (Fig. 7B). Intriguingly, these potential MLL4 direct target genes included *Comp* and *Snorc*, as well as other genes associated with cartilage development and endochondral bone morphogenesis, such as *Matn3*, *Prkg2*, *Scx*, and *Trpv4* (Fig. 7C; Suppl. Fig. 1). Although *Sox9* did not appear in our DEG list, immunofluorescence results clearly indicated reduced SOX9 expression in the *Mll4*-cKO samples (Fig. 5). Incorporating *Sox9* as a “potential” DEG in our analysis revealed that it is targeted by UTX (Fig. 7A, C), strongly suggesting that *Sox9* is a key MLL4 target. Together, these results indicate that MLL4 activates a subset of chondrogenic genes, potentially through the regulation of *Sox9*, in collaboration with UTX. This proposed molecular mechanism is summarized schematically in Fig. 7D.

**Figure 7.**
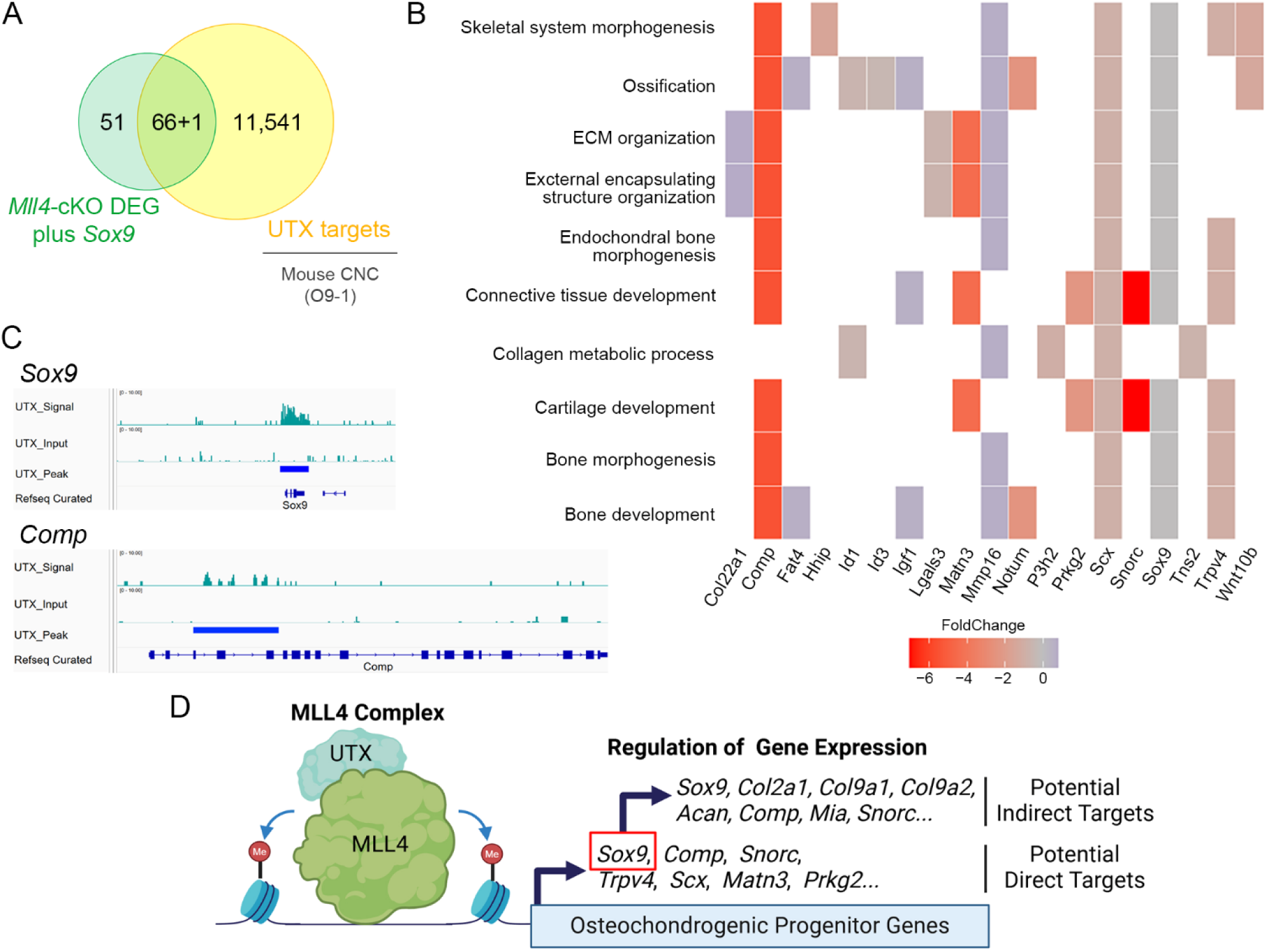
Proposed mechanism by which MLL4 regulates chondrogenic gene expression during midpalatal suture development in postnatal palatogenesis. (A) Venn diagram illustrating the integration of *Mll4*-cKO DEGs (including *Sox9* as a “potential” DEG) and genes identified from the re-analysis of cranial neural crest (CNC) UTX ChIP-seq datasets (Shpargel et al., 2017). (B) GO enrichment analysis of *Mll4*-cKO DEGs overlapping with UTX target genes, highlighting their involvement in various skeletal system-related biological processes, including ‘cartilage development’ and ‘endochondral bone morphogenesis.’ (C) Integrative Genomics Viewer (IGV) browser tracks of representative chondrogenesis-related genes identified by integrating *Mll4*-cKO DEGs and UTX ChIP-seq datasets. The tracks illustrate UTX occupancy at loci of *Sox9* and *Comp*, which are associated with ‘cartilage development’ and ‘endochondral bone morphogenesis.’ (D) Schematic diagram illustrating the deduced molecular model involving MLL4, UTX, and their regulation of potential direct and indirect target genes. MLL4 activates chondrogenic gene expression, potentially through the regulation of *Sox9*, in association with UTX and likely through its role in enhancer priming via by inducing H3K4me1/2 deposition. Created using BioRender.

Another factor contributing to gene expression changes in *Mll4*-deficient midpalaltal sutures is increased cell death. While *Mll4*-cKO mice exhibited a statistically significant increase in TUNEL-positive cells, the percentages of TUNEL-positive nuclei in both control and *Mll4*-cKO mice were relatively low (Fig. 3), indicating that apoptosis is rare in this context. Thus, the widened midpalatal suture phenotype in *Mll4*-cKO mice is likely attributed to deficient development due to impaired chondrogenic gene expression (Fig. 4). Analysis of DEGs for overlap with the Hallmark Apoptosis gene set (Liberzon et al., 2015) identified two downregulated genes, *Igfbp6* and *Lgals3*, but no upregulated genes. SOX9, which is known to regulate apoptosis in neural crest cells (Cheung et al., 2005), may contribute to the observed increase in TUNEL-positive cells (Fig. 3E–G). However, given the low percentage of apoptotic cells, increased apoptotic cells likely reflects an indirect consequence of developmental retardation in the midpalatal suture, rather than being the primary cause of the phenotype.

The chondrogenic retardation observed in our model aligns with previously reported defects in the cartilaginous primordia of the cranial base in neural crest-specific *Mll4* deletion mice and in the growth plates of long bones in heterozygous *Mll4* loss-of-function mice (Fahrner et al., 2019, Shpargel et al., 2020). Also, common downregulated genes were identified when comparing our DEG list with an RNA-seq dataset from earlier palatal shelves (at E14.25) in the neural crest-specific *Mll4* deletion mice (Shpargel et al., 2020), which include *Acan*, *Col2a1*, *Col9a1*, *Col9a2*, *Col9a3*, and *Mia*. These findings suggest a potential consensus regulatory mechanism downstream of MLL4 in different states of cell differentiation at different stages of palatogenesis. Among these genes, *Acan* and *Col2a1* have been identified as causal genes for cleft palate (Xu et al., 2021), highlighting a shared genetic basis for two distinct phenotypes—narrow palate and cleft palate— manifesting under different molecular contexts at their onset. This is consistent with previous studies showing that high-arched, narrow palate and cleft palate may share common genetic causes (Conley et al., 2016, Tabler et al., 2013). However, it remains unclear why bilateral palatal shelves fail to fuse only in the neural crest-specific *Mll4* deletion mice (resulting in cleft palate) but not in our palate mesenchyme-specific *Mll4*-cKO mice. This may reflect an earlier requirement for MLL4 during the migration and differentiation of neural crest cells prior to their incorporation into the palate mesenchyme, potentially explaining the cleft palate phenotype in neural crest-specific *Mll4* mutants.

For future studies, it will be important to investigate whether the H3K4 methyltransferase activity of MLL4 is specifically required for its role in postnatal palatogenesis. Recent findings (Xie et al., 2023) suggest that MLL4 has both H3K4 methyltransferase-dependent and -independent roles in development and differentiation. Notably, while H3K4 methylation appears largely dispensable for enhancer activation during embryonic stem cell differentiation, it is crucial for lineage-selective processes in early embryonic development. Given that MLL4 predominantly mediates the deposition of H3K4 mono- and di-methylation (H3K4me1/2) (Lee et al., 2013, Ang et al., 2016, Wang et al., 2017), its enhancer-related functions may be particularly relevant during postnatal palatogenesis. Exploring these distinct mechanisms in the context of enhancer priming and tissue-specific development may provide valuable insights into the regulatory functions of MLL4.

### MLL4 and the midpalatal suture: a central growth site and mechanosensitive regulator of craniofacial development

The midpalatal suture has long been regarded as a critical skeletal structure that is anatomically situated at the very center of the craniofacial complex, playing a significant role in the development and growth of the palate and the maxillary jaw (Latham, 1971, Li et al., 2015, Rice, 2008). Therefore, it is intriguing to discover that the two major phenotypes observed in the *Mll4*-cKO mice were a growth deficit in the palate and developmental retardation of the midpalatal suture. While the causal relationship between midpalatal suture developmental retardation and lateral growth deficit remains to be tested in future studies, our findings invite the possibility that MLL4 regulates growth of the palate and the midface by modulating midpalatal suture. This highlights a functional link between midpalatal suture development and palate growth and supports the potential central role of the midpalatal suture as a growth site driving craniofacial development.

In addition to serving as a growth site, the midpalatal suture functions as a mechanosensitive structure that responds to environmental forces, such as suckling, mastication, and skeletal growth (Roth et al., 2022, Li et al., 2016). For example, in orthodontic dentistry, rapid palatal expansion procedures apply mechanical force to the maxillary jaw, widening the midpalatal suture and inducing tensile forces that promote new bone formation, thereby increasing maxillary width. These responses have been partially attributed to *Piezo2*-positive chondrogenic mesenchyme cells in the midpalatal suture (Gao et al., 2022). Therefore, it would be interesting to test whether *Mll4* deficiency impacts the mechanosensory properties of the midpalatal suture, possibly through *Piezo2*-positive chondrogenic mesenchyme cells. This can be explored by simulating mechanical stress, such as performing a palatal expansion procedure in *Mll4*-cKO mice (Hou et al., 2007, Katebi et al., 2012). Such studies would provide valuable insights into the role of MLL4 in craniofacial development and the biological responses of the midpalatal suture to mechanical cues.

## Data Availability Statement

Raw expression data results from the RNA-seq are included in the Supplementary Material, and the original RNA-seq data files were deposited in the NCBI GEO database under the accession number PRJNA1148613.

## Ethics Statement

The animal study was reviewed and approved by Institutional Animal Care and Use Committee (IACUC) at the University at Buffalo.

## Author Contributions

J-ML acquired funding, conceptualized, designed, administered, and supervised the research, performed experiments, acquired data, conducted data analysis and visualization, wrote and edited the manuscript. HJ performed experiments, acquired data, conducted data analysis and visualization, and edited the manuscript. BdPMP performed experiments, acquired data, conducted data analysis and visualization, and edited the manuscript. YP conducted data analysis and visualization and edited the manuscript. QT conducted re-analysis and visualization and edited the manuscript. SJ provided resources. S-KL acquired funding for the project, provided resources, and edited the manuscript. JWL acquired funding for the project, provided resources, and edited the manuscript. H-JEK acquired funding, conceptualized, designed, administered, and supervised the research, performed experiments, acquired data, conducted data analysis and visualization, wrote and edited the manuscript. All authors contributed to and approved the publication of the manuscript.

## Acknowledgements

We thank Andrew D. McCall at the Optical Imaging and Analysis Facility (OIAF), School of Dental Medicine, The State University of New York at Buffalo, for assistance in acquiring micro-CT data and gross, histology, and stereo immunofluorescence images, which were all conducted at the OIAF. We thank Yang Chai at the University of Southern California for providing the *Osr2-Cre* mouse strain. We thank Eun-Myung Mary Song and Monique M. Kapur-Mauleon for assistance with experiments. We would also like to thank the members of the Kwon lab for their helpful comments and feedback.

---

**Supplemental Figure 1.**
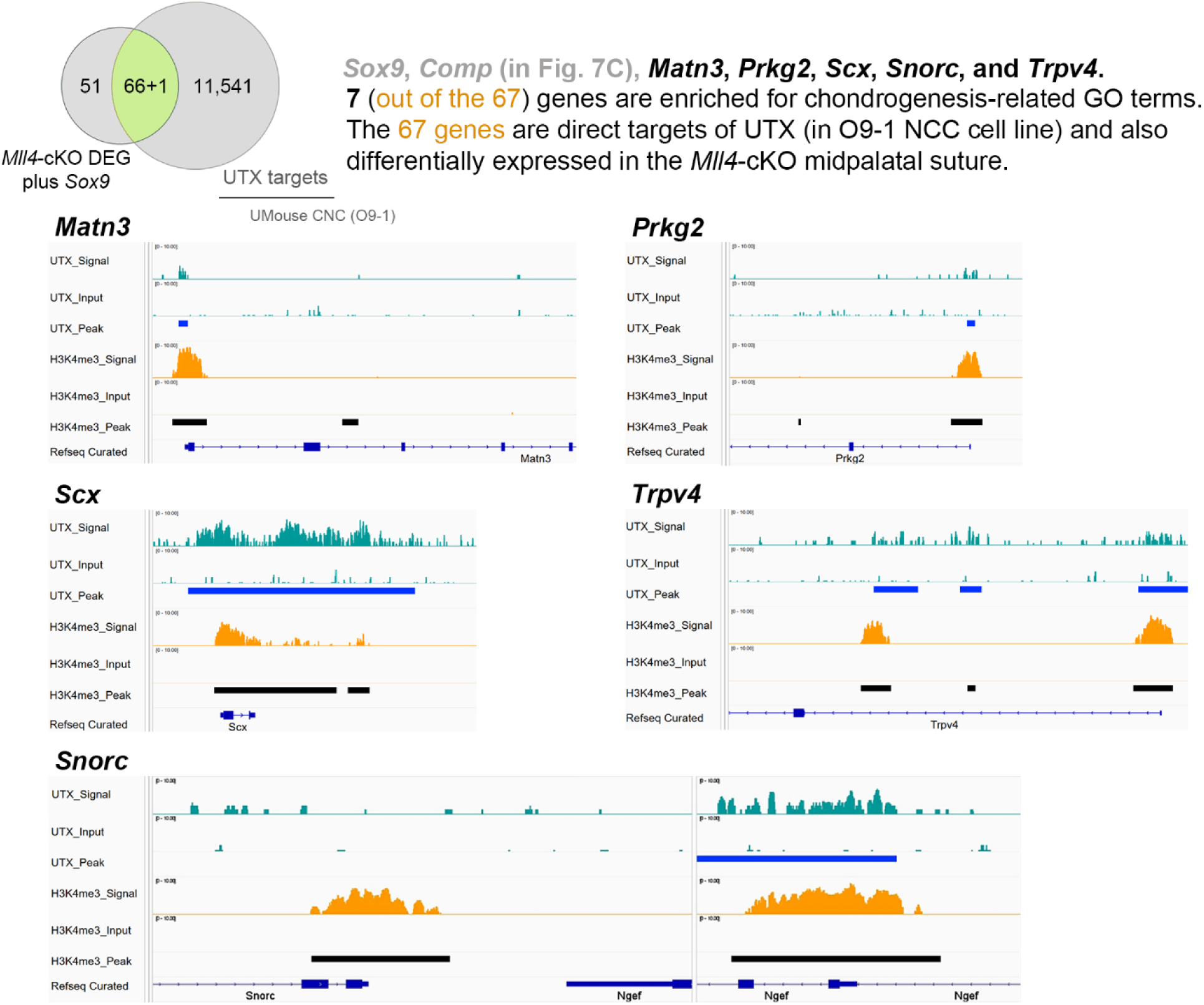
Integrative Genomics Viewer (IGV) browser tracks of UTX ChIP-seq results. Representative chondrogenesis-related genes identified by integrating *Mll4*-cKO midpalatal suture DEGs with O9-1 cranial neural crest UTX ChIP-seq datasets (Shpargel et al., 2020). The tracks illustrate UTX occupancy at loci of genes such as *Matn3*, *Prkg2*, *Scx*, *Snorc*, and *Trpv4* which are associated with ‘cartilage development’ and ‘endochondral bone morphogenesis,’ suggesting that MLL4 functions in coordination with UTX to regulate these genes, likely through enhancer priming via H3K4me1/2 deposition.

