## Supplementary figures and images for "MLL4 regulates postnatal palate growth and midpalatal suture development"

### Supplemental Figure 1

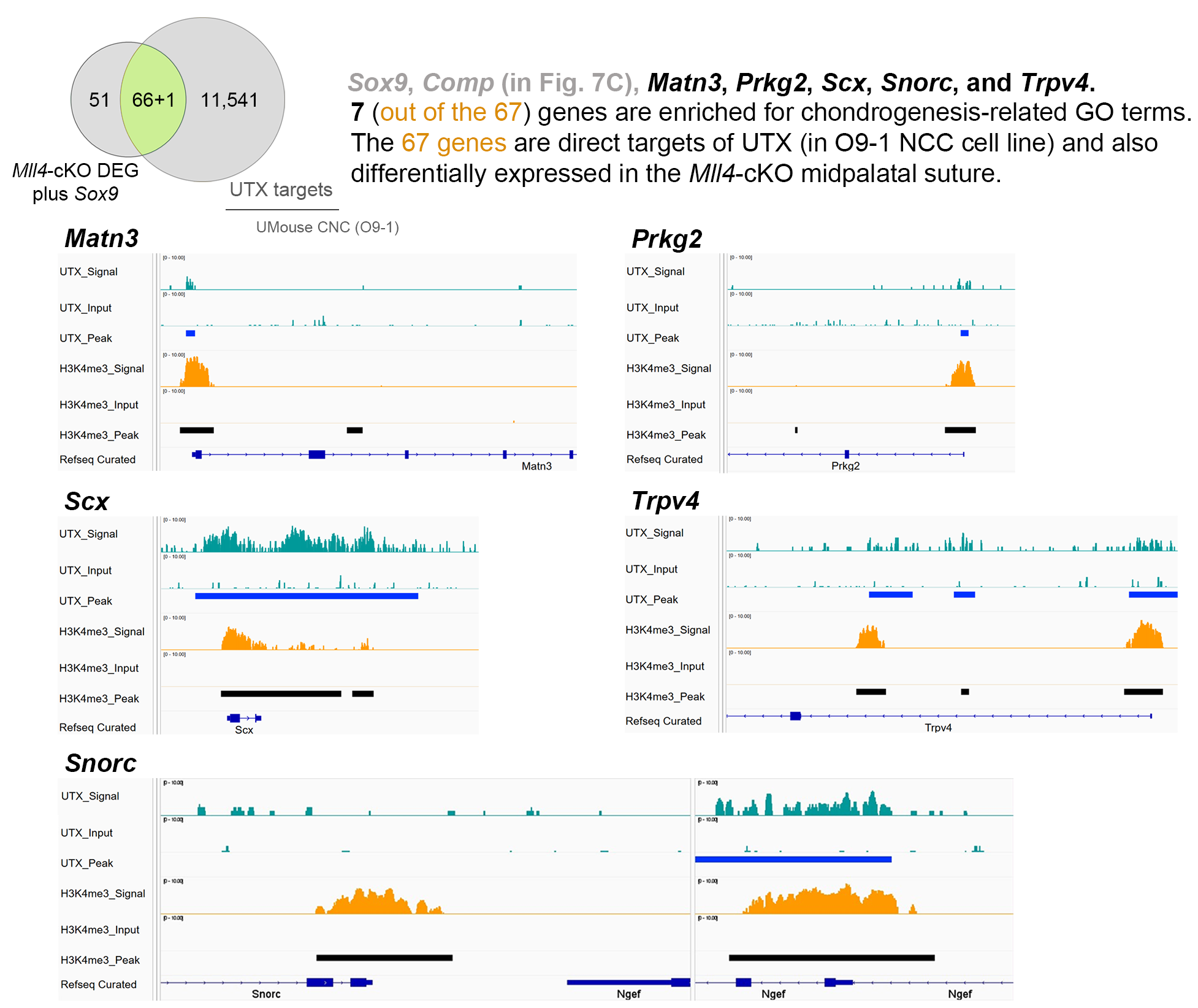
